# Quantification of anatomical changes in young grapevine wood over time and in response to *Neofusicoccum parvum* with image processing

**DOI:** 10.64898/2026.03.20.713180

**Authors:** Célia Perrin, Jean-Baptiste Courbot, Yann Leva, Romain J. G. Pierron

## Abstract

Grapevine Trunk diseases (GTDs) represent a major threat for the wine industry. Despite several break-through, their etiology remains unclear and no curative treatment is currently available. Wood anatomy and water transport contribute to the symptoms of young plant decline. This study investigates wood anatomical alterations in two Alsatian grapevine cultivars presenting different susceptibility to GTDs, focusing on wood structure over six months of vegetative growth and in response to infection. Using a validated FasGa staining protocol, wood sections from transverse, tangential, and radial directions were stained to differentiate lignified and cellulosic tissues. Microscopic analysis was performed at x4, x10, and x40 magnifications, yielding a dataset of 4771 images. To support this high-throughput quantitative analysis of microscopy images, a computational model was developed, enabling reliable and efficient assessment of anatomical traits. Pre-established woody tissues presented higher xylem vessels diameter in Gewurztraminer than Riesling, with a dorsoventral arrangement whereas the number of vessels remained the same all over the cross section. No significant anatomical changes were observed in established woody tissues, whereas newly formed xylem anatomy showed a possible rearrangement during infection, especially in Gewurztraminer cultivar. Furthermore, colorimetric analysis quantified the lignification of woody tissues in response to wounding damage compared to untreated plants. While definitive conclusions remain limited due to the experimental timeframe and sample variability, the findings highlight the need for longer-term studies and broader cultivar evaluation. Code and microscopy images have been made publicly available, providing a scalable digital tool for future research in plant vascular systems.

## 1. Introduction

Grapevine Trunk Diseases (GTDs) represent a worldwide threat for the wine industry and important economic losses estimated in excess of 1 billion euros only regarding plant death replacement [1]. In France, 12 % of the vineyard is unproductive because of these diseases and regions of Alsace and Upper Rhine, which play an important role in wine industry, are not spared [2]. GTDs include Esca, Eutypiose or Black Dead Arm (BDA), symptomatic plants generally presenting a loss of vigor and chronic decline which often lead to premature death of plants, also referred to as young grapevine decline [3–5]. Since the ban of sodium arsenite in 2001 in France because of its risk on human health and environment [6], there is currently no efficient and long-term controlling tool available [7].

A cocktail of fungal species colonizing the woody tissues of grapevine trunks causes wood degradation which is associated with characteristic foliar symptoms. The diversity of foliar symptoms expression makes the difference between Esca, Eutypiose or BDA. The link between fungal pathogens inhabiting the xylem tissues, wood degradation and foliar symptoms expression is not clearly established, excepted for Eutypiose for which foliar symptoms have been reproduced in laboratory conditions [8,9]. Regarding Esca and Botryospheria dieback, only two papers have reported partial reproduction of foliar symptoms [10,11]. Pathogens gradually degrade wood tissues with the formation of internal necrosis and obstruction of vessels. These diseases keep progressing, probably supported by climate change and its resulting abiotic stresses. These conditions are favorable to the vascular development of a complex of pathogenic fungi which will degrade the wood and cause foliar symptoms [12]. Some grape varieties are particularly susceptible to these diseases without elucidating clearly the mechanisms of tolerance or sensitivity involved. In Alsace, Riesling presents a higher tolerance than Gewurztraminer when averaging symptom occurrence from 2003 to 2022 [13]. If environment or cultural factors can influence the dynamics of expression of these diseases, foliar symptoms expression in vineyard differs among cultivars, which could suggest that there are very specific physiological or anatomical determinants enabling the development of these diseases [14]. Gastou and colleagues compared the influence of water use strategy, wood anatomy, metabolome and microbiome among 46 *Vitis vinifera* genotypes regarding foliar symptom expression of Esca. Interestingly cultivar differences in water use and conservation strategy seems to prevail on wood anatomy when correlating the data of leaf gas exchange and microscopic observation of woody tissues with foliar symptoms expression. Some metabolic changes were recorded but no differences in microbial communities when comparing symptomatic and asymptomatic plants [15].

Wood properties condition the mechanical support, the sol-plant-atmosphere continuum, biochemical compound storage involved in plant development, reproduction, and defense [16]. Wood anatomy should thus be considered as a major trait when investigating plant resilience toward environmental stress [17,18]. The Compartmentalization Of Decay in Trees (CODIT) model proposed by Shigo and Marx [19] theorizes a strategy to separate the cambium from trunk infection sites, favoring long term survival in trees. Four compartmentalization walls are associated with trunk defense, three walls being constitutive defenses. Interestingly, the fourth originates from the cambium present at the early events of the infection, suggesting the existence of induced defense in the trunk tissues. If the xylem tissue is often considered to be made of dead cells because of the xylem vessels, it is surrounded by pluripotent cambial cells, and adjacent parenchyma also contains active living cells especially in sapwood [20,21]. Induced plant defenses imply a finely tuned perception of fungal pathogens by living xylem cells, as described by the Zig-Zag model [22]. Several studies highlighted the existence of stress perception and defense induction in woody tissues [23]. In *Populus* sp., single-nucleus transcriptomics highlighted auxin-driven mechanisms involved in response to severe drought stress. Drought perception induces early vessel differentiation, leading to the formation of secondary xylem with narrower vessels in higher numbers [24]. It is also the case in *Ulmus minor*, which presents important transcriptomic changes in response to *Ophiostoma novo-ulmi* [25].

Compartmentalization walls from the CODIT model are also observable in grapevine, such as circular compartments to contain white rot associated with *Fomitiporia mediterranea*, or V-shaped black necrosis associated with *Botryosphaeriaceae* species [26]. Several GTD-associated fungi can cause a wood degradation when inoculated in laboratory conditions such as *Phaeomoniella chlamydospora* [27] or *Neofusicoccum parvum* [28]. The anatomical structure of grapevine wood, particularly the architecture and functionality of the vascular system, may play a key role in the response of grapevine cultivar to fungal infections. Pathogen infections or unfavorable climatic conditions can alter the development of its anatomical structure. Rizzoli and colleagues observed that grapevines affected by “Flavescence dorée” showed reduced xylem and phloem development, particularly noticeable in the size of the growth ring boundaries [29]. Similarly, under drought conditions, the plant undergoes physiological adjustments to make more efficient use of available water resources, which also results in reduced wood structure development and smaller vessel sizes [29,30]. Several studies have shown that vessel sizes may be associated with increased susceptibility of the grape cultivar to pathogens. Larger vessels may facilitate the movement of pathogens through the vascular system, promoting their spread and intensifying symptoms expression [31]. Although this anatomical trait improves water transport efficiency, it also increases vulnerability to infection by fungi such as *Phaemoniella chlamydospora* [32]. However, wide vessels are also sites of active defense responses, including the formation of tyloses and gels that compartmentalize the pathogen and help to limit its progression [33]. This trade-off between hydraulic efficiency and pathogen sensitivity is a key factor to consider when investigating the differences in cultivar responses to vascular pathogens. Gastou and colleagues present the importance of xylem anatomy associated with Esca foliar symptoms and suggest the importance of newly formed xylem tissues regarding the resilience of grapevine toward GTDs [15].

To our knowledge, there is currently no data measuring temporal variation in wood anatomy of young grapevine plants. In addition, image analysis in microscopy can be time-consuming and tools adapted to grapevine wood anatomical studies are also missing for our research community. This study aims to monitor anatomical variations in the one-year-old wood of grapevine cultivars Riesling and Gewurztraminer, healthy or inoculated with the fungal pathogen *N. parvum* strain Bt67. A particular focus on the distribution, size, and density of xylem vessels and newly formed xylem tissues was made. The present work is associated with a data set of 4771 images of grapevine wood, together with an automated image processing pipeline available online.

## 2. Materials and Methods

### 2.1 Biological material

One-year-old canes of *Vitis vinifera* L. cv. Gewurztraminer and cv. Riesling were harvested in December 2023 in the INRAE vineyard (Colmar, Alsace, France). These two grapevine cultivars were selected for their high susceptibility to GTDs in Alsace. Canes were divided into cuttings containing two dormant buds and were waxed after a hot water treatment consisting of a 45-minutes bath at 50°C. Cuttings were planted in moistened rock wool in plant growth chamber (16/8 photoperiod, 80% relative humidity, 26°C) until budding and rooting.

In the meantime, the fungus *Neofusicoccum parvum* isolate Bt67 (*Np*Bt67) was cultured in potato-dextrose agar (PDA) at 27°C, in the dark and subcultured every 10 days. This strain was obtained from the Higher Institute of Agronomy in Lisboa, Portugal.

### 2.2 Plant treatment, inoculation and sampling

One month after budding, when roots were well developed, 105 grapevine cuttings per cultivar were transplanted into one-liter pots containing natural pozzolana (7/14mm, Sorexto, France) at the bottom to prevent root hypoxia, and horticultural soil (TH110/36 supplemented with 7-4.5-12 NPK, Dumona, France). All pots were placed uniformly in a growth chamber and randomly selected for the experiment when plants had six fully developed leaves (c.a. two months after rooting). All plants were maintained under the same conditions in a plant growth chamber (16/8 photoperiod, 65% relative humidity, 25°C) and well-watered twice per week, for a maximum of 6 months from April to September 2024. To avoid a potential impact of potting stress, treatments were applied two weeks after potting as follows, starting in March 2024 (Fig.1):

*(I) Anatomy.* 60 plants per cultivar for the study of grapevine wood anatomy from 1 to 6 months post potting stress.
*(II) Response to infection.* 30 plants per cultivar for the analysis of wood anatomy in response to infection by *Np*Bt67. Three conditions will be referred as control, mock, and infected, respectively and contained 10 plants each. To proceed, plants were wounded using a drill (Black and Decker, USA) with a 5-mm diameter bit. The hole was made at the middle of an internode for the treated plants and inoculated especially with a 5 mm diameter plug of PDA alone (mock) or a plug of 10-days old *Np*Bt67 subculture (infected). Wounds were finally sealed with a piece of parafilm. Control plant remained undamaged.

**Figure 1:**
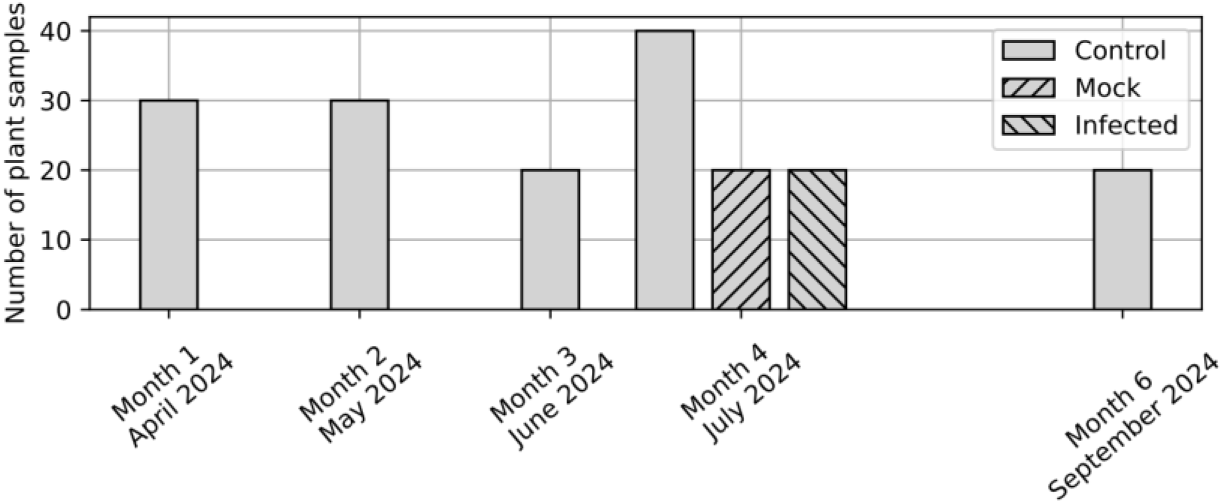
Number of plants sampled at each time point of the study for the two grapevine cultivars, Gewurztraminer and Riesling. Note that the same number of plants was assessed in both cultivars.

For each sampling, wood was cross-sectioned with a circular saw (Proxxon KS230, Germany) into 1.5 cm long sections. Regarding the section on wood anatomy over time (I), the trunk cuttings of 10 to 15 plants per cultivar were collected monthly for 6 months starting in April 2024. Initially, 15 plants were intended to be sampled at each time point; however, some plants died over the course of the experiment. Consequently, the remaining plants were distributed among the different time points and sampled as summarized in Fig.1. Plants inoculated with PDA or fungal mycelium (II), were sampled in sections made 1 cm above and below the inoculation point. These fragments were stored at 4°C in 70% ethanol in sterile 15 mL centrifuge tubes before processing the samples for observations. Undamaged control plants from (II) were grouped with healthy individuals in (I) at the month 4 sampling point because plants were grown under the same conditions. Therefore, they were combined with the month 4 dataset for analysis to increase the robustness of our study.

### 2.3 Preparation and coloration for optic microscopy

For each sample, wood sections were sliced in the three cutting directions (transversal, tangential, and radial). For each direction, several 18 µm thick wood sections were made using a lab-microtome (WSL, Switzerland). All samples were stained with the FasGa dye adapted from Tolivia and Tolivia’s [34] protocol. This dye is composed of a mixture of 0.08% Safranin O (Carl Roth), 0.14% Alcian Blue 8GX (Sigma Aldrich), 1.3% acetic acid (Carl Roth) and 38% glycerol (Fisher Chemical) in distilled water. The dye was selected for its ability to specifically differentiate lignified structures, which appear red, from cellulosic structures, which appear blue. The slides were incubated for 8 hours in FasGa dye diluted in distilled water (1:7, v/v) and rinsed for one minute in three successive baths of 100% ethanol. Then, they were fixed immediately after rinsing with Euparal glue (Carl Roth) and left to dry for 1 week before starting the observation.

### 2.4 Microscopic and macroscopic observations

All observations, on the three cutting directions, were carried out using an optical microscope (Motic Optic BA210E, China) and done at objectives x4, x10 and x40. For each observation direction, 4 to 6 microscopy images were captured. Regarding the cross-section, an additional observation was made with a macroscope (Axio Zoom V16 ZEISS, Germany) using x8 or x10 objectives, depending on the cross-section size, to obtain a wide-field image integrating the entire section.

Overall, these observations form a large database of 4771 images, covering all the variations described above: two cultivars, five-time steps, three infection conditions at month 4, 10 to 15 samples, that were each observed in microscopy (3 directions) and with a macroscope. A sample of the database is given in Fig.2 and 3. This database can benefit the research community, and is publicly available online at https://doi.org/10.5281/zenodo.18850060 [35].

**Figure 2:**
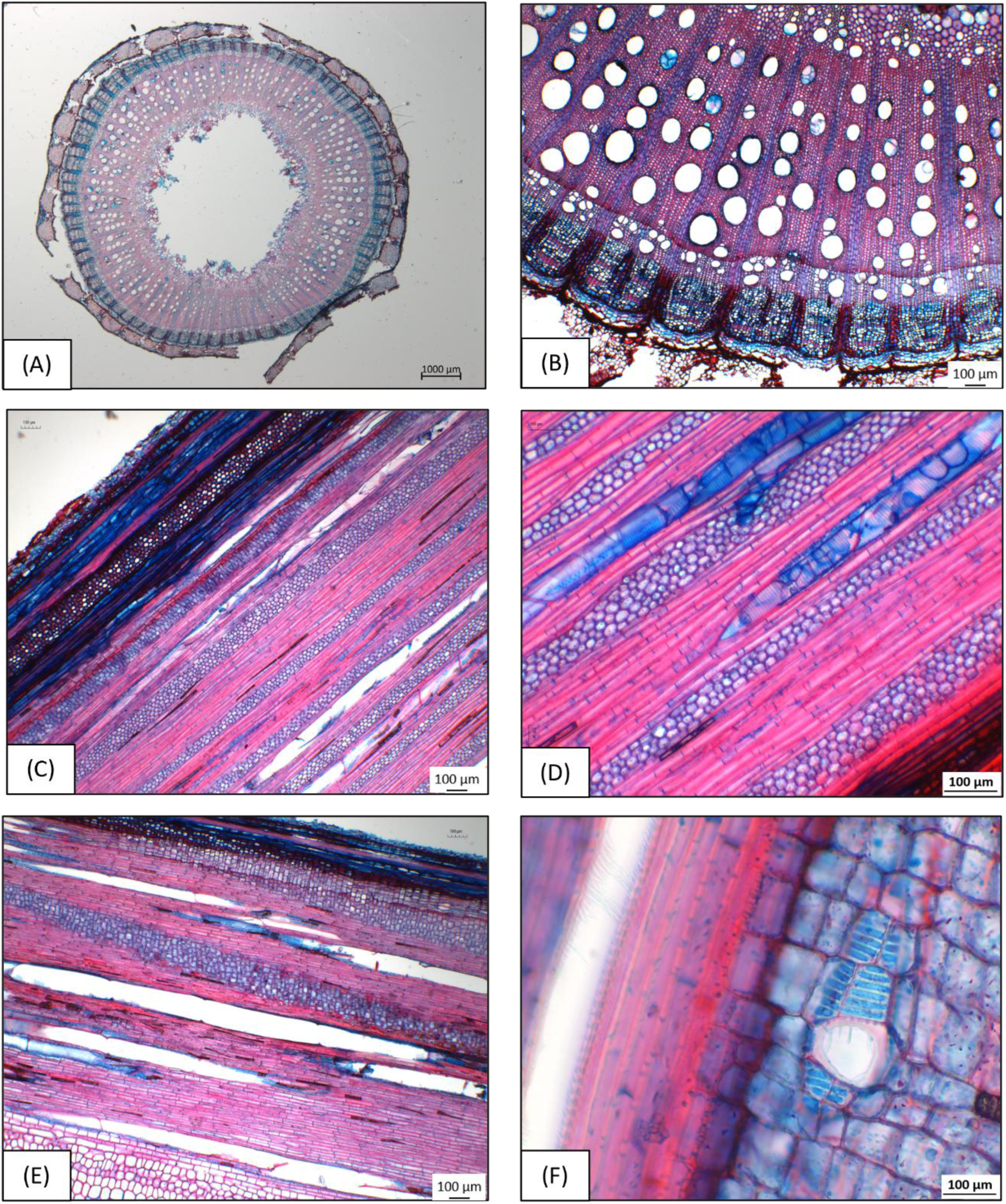
Typical anatomical observations of the Gewurztraminer cultivar in the three cutting directions. Each wood sample was examined in the three anatomical directions. Macroscopic observations were conducted on transverse sections to obtain an overall view of the sample using a x10 objective (A). Microscopic observations were performed using an optical microscope at different magnifications (x4, x10 and x40): in transverse section (B, x4), in tangential section (C, x4 and D, x10) and in radial section (E, x4 and F, x40), allowing detailed visualization of the different wood compartments.

**Figure 3:**
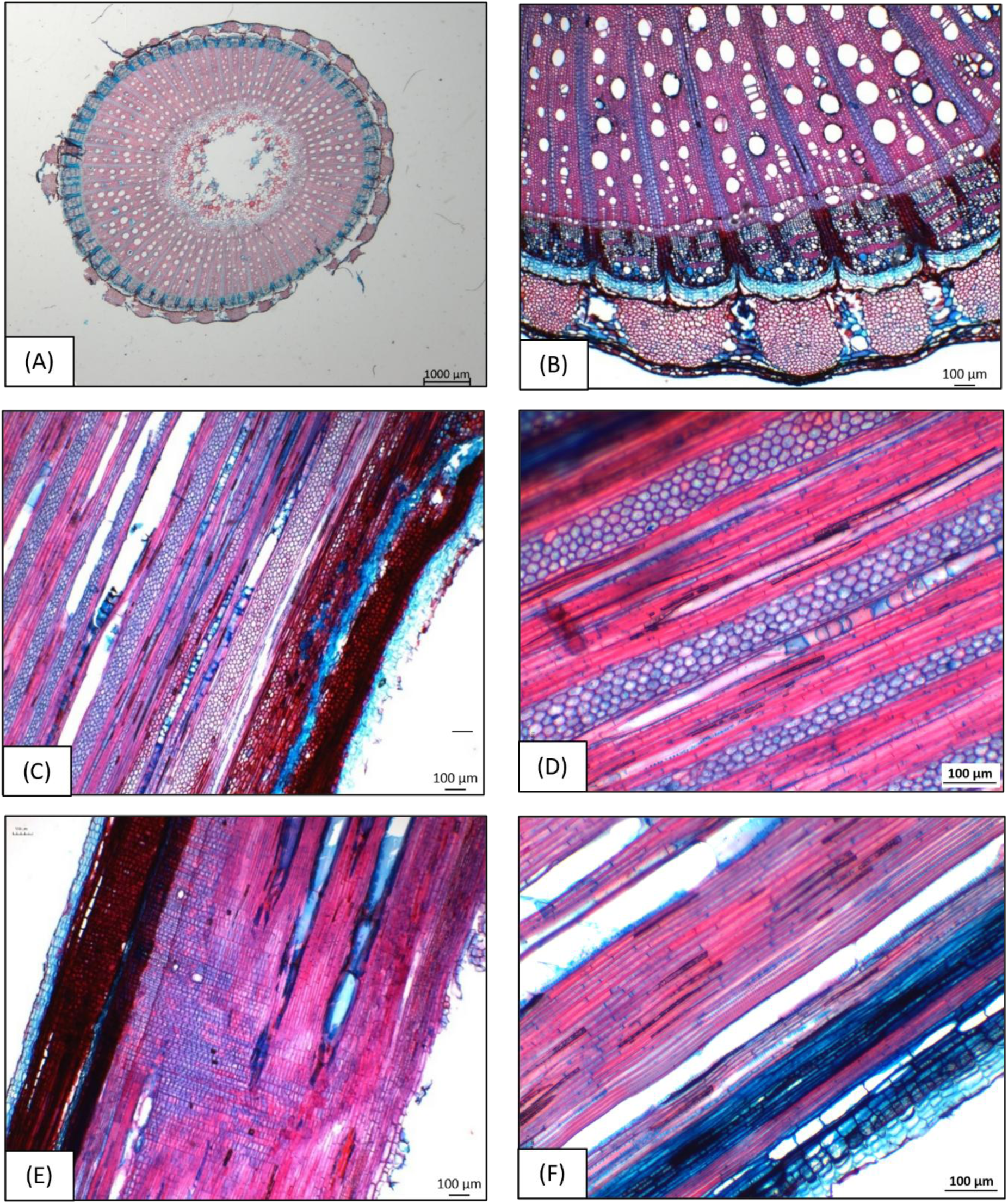
Typical anatomical observations of the Riesling cultivar in the three cutting directions. Each wood sample was examined in the three anatomical directions. Macroscopic observations were performed only on the transverse section to obtain an overall view of the sample using a x10 objective (A). Microscopic observations were carried out using an optical microscope at different magnifications (x4, x10, and x40): in transverse section (B, x4), in tangential section (C, x4 and D, x10) and in radial section (E, x4 and F, x10).

### 2.5 Image processing

Given the size of the image database, automated image processing tools were developed in order to quantify several morphological properties of the wood. The Python source code to reproduce the quantification procedure is available online at <u>github.com/courbot/vineside</u>.

#### 2.5.1 Xylem vessels morphology

The first studied aspect is the morphology of vessels within the xylem. Macroscope cross-section images were used, enabling a sufficient resolution to map vessels individually. This image segmentation task was performed automatically, following the procedure described below:

a. Automatic segmentation of the image using Meta’s Segment Anything Model (SAM) [36]. This step allows discriminating different regions of the image along their shape and texture.
b. Object removal depending on their size: x10 images enable the determination of minimum and maximum vessel size in xylem for each cultivar. These values were used to reject outliers (too small or too large shapes) from our sample.
c. Object removal depending on their closeness: objects too far from the center of the image are likely to be located outside the wood, and thus of not being a xylem vessel. They were also removed.
d. Finally, the automatic segmentation highlighted marrow cells that are irrelevant to the study;they were removed based on a vicinity criterion. The measure of the width of marrow cell walls also enabled to reject objects too close from another.

This enables to distinguish individual vessels and to compute for each one their surface, position, and eccentricity. The study especially focuses on locating the vessels in polar coordinates, that is, their distance and orientation with respect to the center of the image (Fig.4). This will ensure the measurement of their spatial repartition, especially the morphological properties of the vessels with respect to their coordinates.

**Figure 4:**
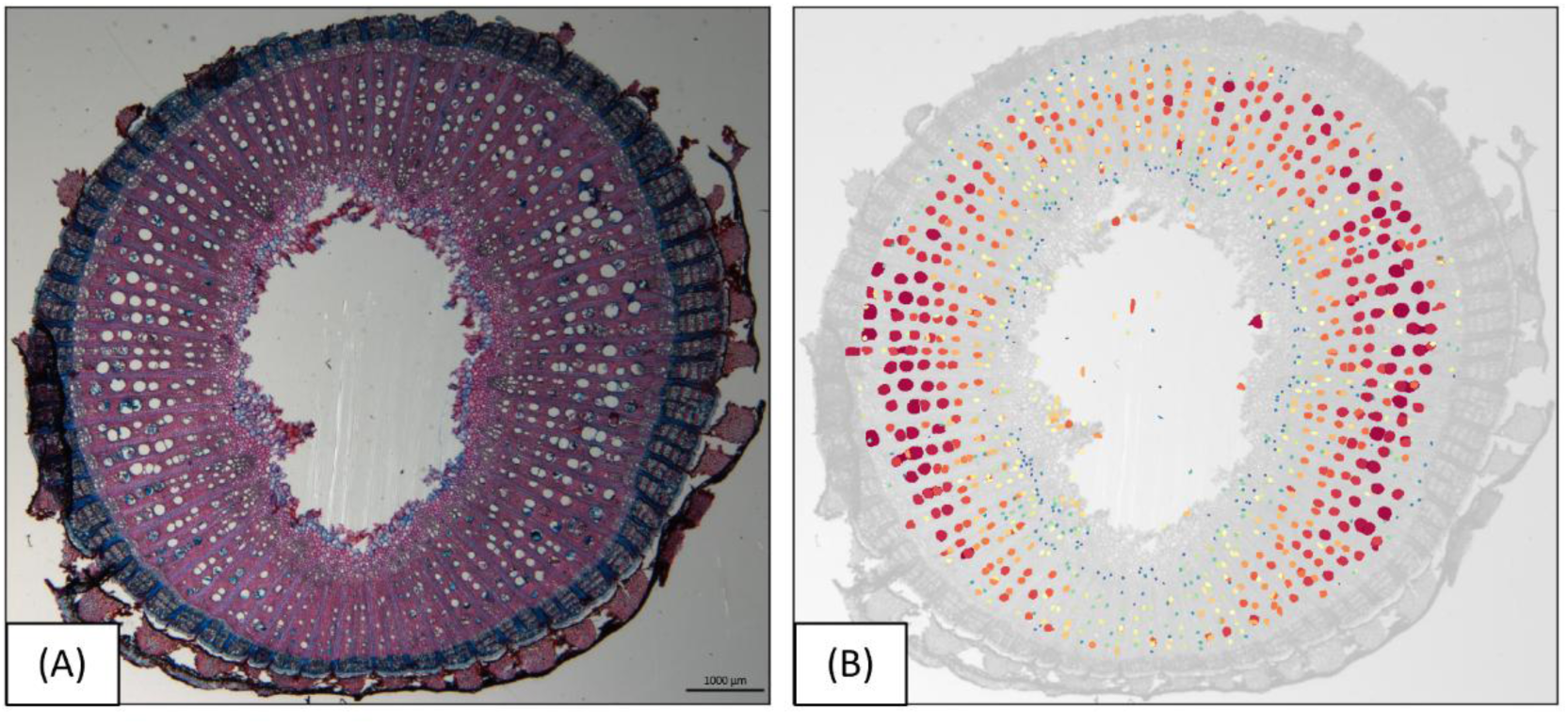
Example of a macroscope cross-section (A) and resulting automatic xylem vessel detection (B). Colors distinguish between different vessels and are ordered by size.

#### 2.5.2 New xylem morphology

Macroscope images also allow quantifying the new xylem, that is, the part of the xylem that has been formed over the duration of the experiment. As it is difficult to automatically identify, manual segmentations were performed, yielding examples such as those presented in Fig.5. Then, two morphological features of the new xylem are observed:

a. *New xylem thickness.* This can be quantified through the computation of two quantities: the distance transform of the binary image, and its skeletonization [37]. This yields a skeleton of the new xylem mask and maps the distance to its border, thus enabling to quantify for each cross-section an average new xylem thickness. The overall procedure is summarized in Fig.5.
b. *Morphology of new xylem vessel*. By taking the intersection of the new xylem mask with the automatic vessel segmentation presented in Fig.4, it is possible to separate vessels within and outside the new xylem. This enables, in turn, quantifying the new xylem vessel area and count, depending on the experimental conditions.

**Figure 5:**
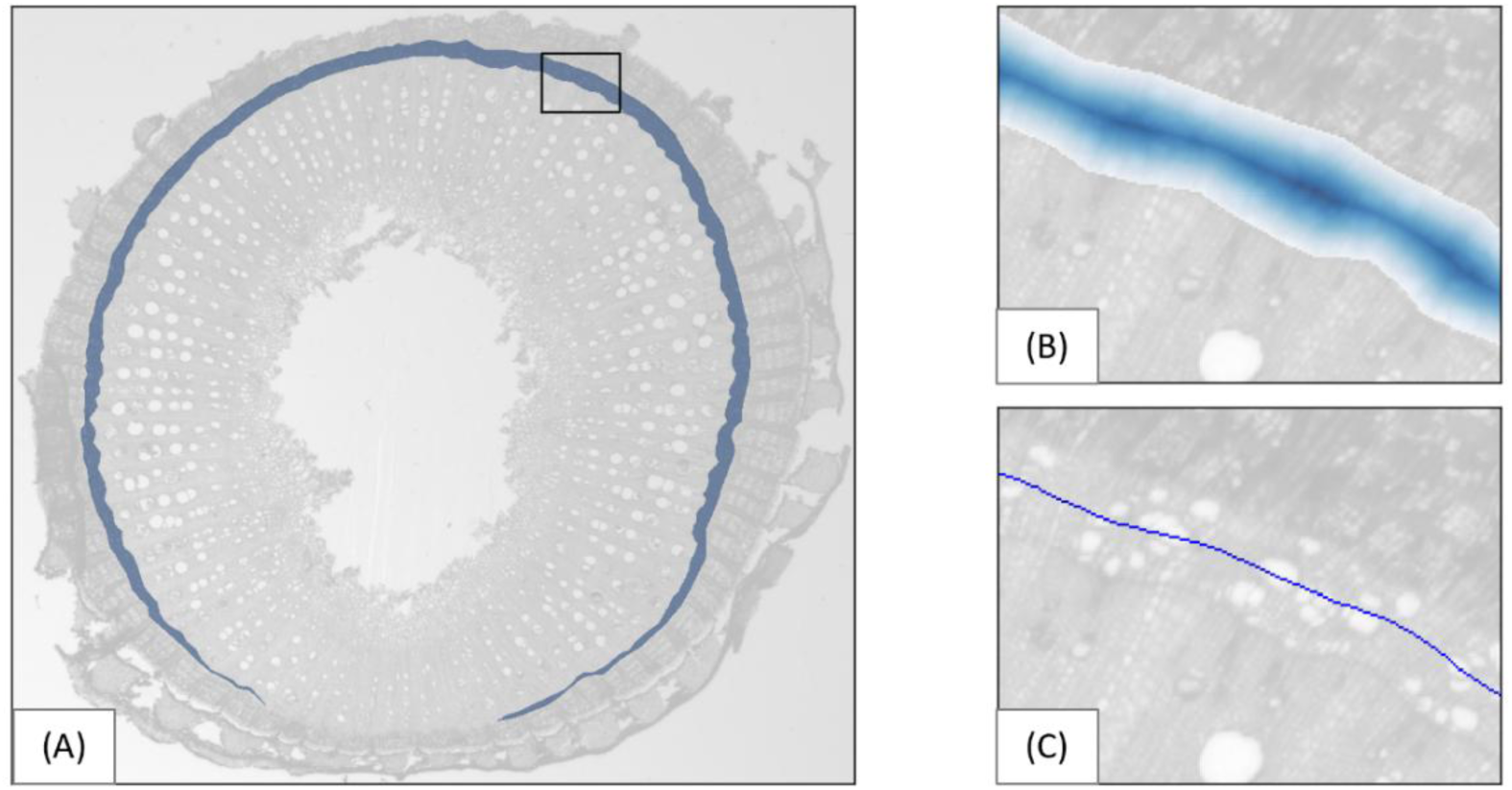
New xylem manual segmentation (A) and computation of its thickness using the distance transform (B: darker shade represents higher distance to the border) and skeletonization (C).

#### 2.5.3 Colorimetry

The FasGa dye is designed to enhance lignified (red) and cellulosic (blue) structures, thus enabling the quantification of the color balance within the image. To do so, the microscopic x4 images of cross-section slices were investigated. Meaningful clusters of pixels were identified using the SLIC over-segmentation technique [38], which gathers pixels based on their distance in the image and their similarity in color. Then, clusters presenting similar levels in the (R, G, B) channels were removed, in order to exclude from the study, the regions having an average color close to black, gray, or white. The procedure is depicted in Fig.6 and yields an average color per image, that is in turn averaged between observations of the same plant, in order to compute an average color per plant sample. To avoid accounting for brightness, the red and blue colors are also normalized by the pixel intensity.

**Figure 6:**
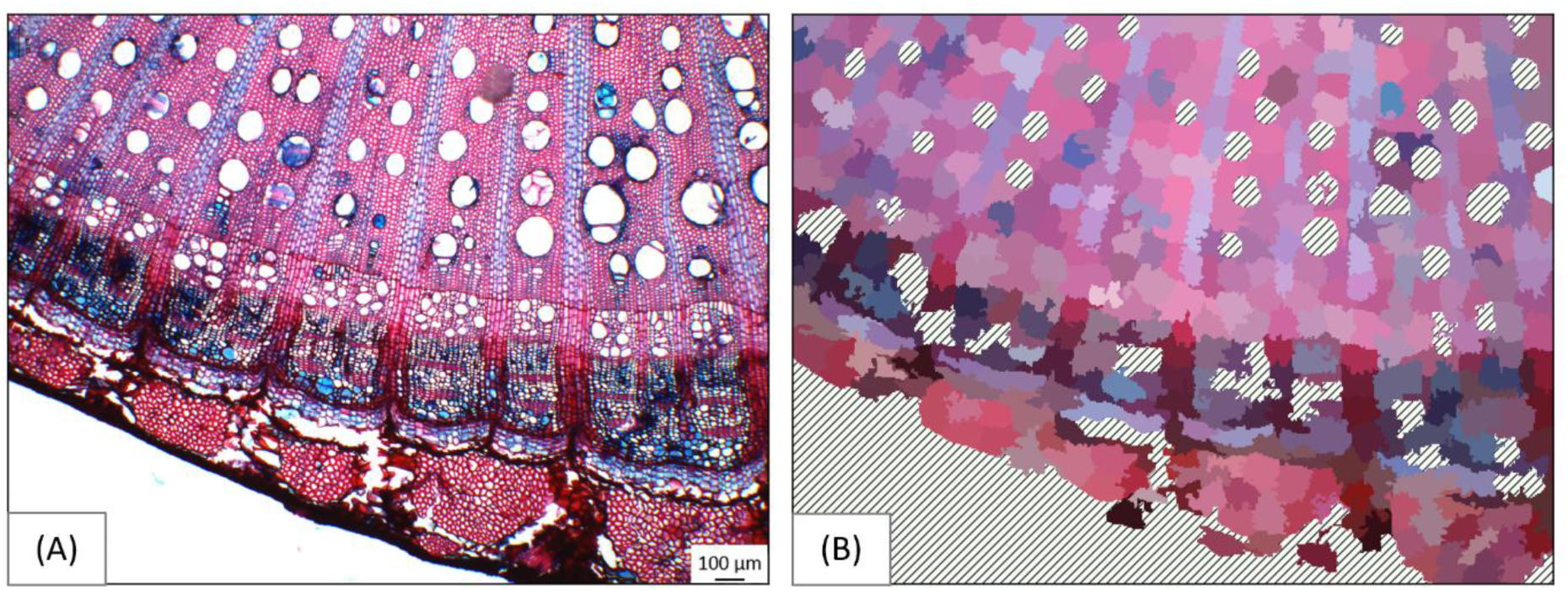
Colorimetry procedure: from the microscopic image (A), we cluster pixels together (polar patches in (B)) while excluding regions without contrast between colors (hatched in (B)).

#### 2.5.4 Phloem thickness

The thickness of the phloem is also measured from radial slices observed in x4 microscopy. In these images, the phloem is the darker region. To isolate it, SLIC clustering is used to regroup pixels by distance and color. Then, thresholding the result yields a binary mask whose largest element is the phloem. Finally, its thickness was measured according to the skeletonization procedure depicted for the new xylem thickness study. This method, depicted in Fig.7, yields a median phloem thickness for each image, that is averaged within each plant so that there is a measure of phloem thickness per plant sample. Results were manually checked, as phloem is not always present in the samples, and, when present, may be observed in several distinct layers. Thus, only phloem in one piece, as in Fig.7, is accounted for in this study.

**Figure 7:**
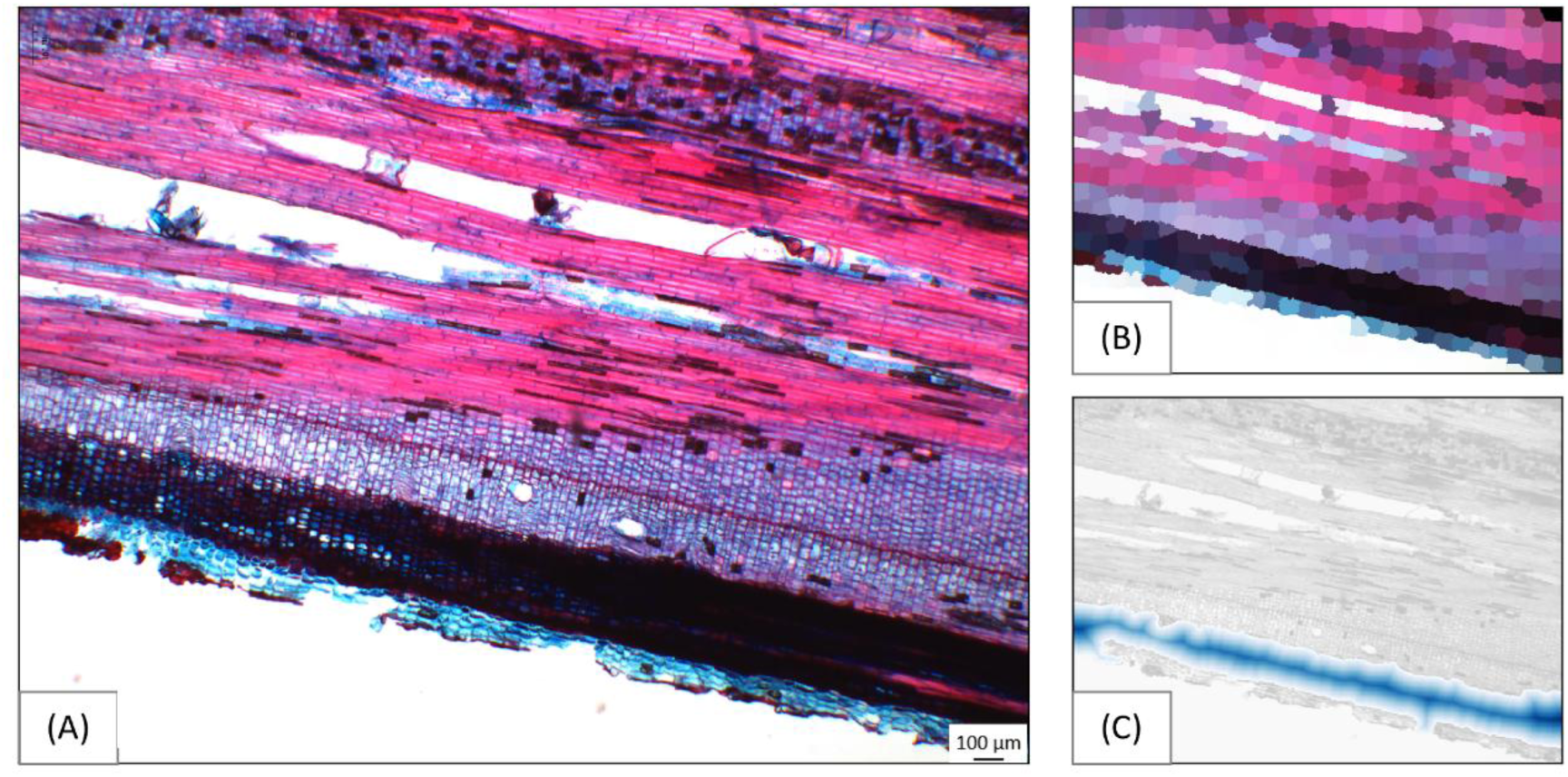
Illustration of the phloem thickness measurement on x4 microscopy images of radial wood slice (A). B: SLIC over-segmentation to cluster pixels by location and color, allowing the detection of the largest darker shape. Then, median distances are computed in the same way as for the new xylem thickness (C, as in Fig.5).

Summarizing, the following parameters were measured:

a. Individual vessel morphology: positions in polar coordinates, surfaces, density, and eccentricity. Here, a sample individual is a xylem vessel.
b. New xylem average thickness, with one measure per plant.
c. Colorimetry in red and blue channels, with a pair of (red, blue) measures per plant.
d. Phloem thickness, with one measure per plant.

Note that all image processing tools mentioned here are not expected to be perfect and may yield spurious results. To mitigate this, the results were manually checked, and only a small amount of erroneous output was found. Thus, their impact on the results presented afterward is expected to be negligible.

### 2.6 Statistical methodology

This section addresses how sample differences were established in the various conditions (time steps, infection, etc.). To do so, as samples are statistically small (10 to 15 samples), a Kruskal-Wallis ranking test was adopted for comparison. The null hypothesis H_0_ for this test is that all considered samples come from the same distribution.

The temporal evolution of features is analyzed using a linear regression. To assess the statistical significance of the observed trend, a Wald test is performed, with the null hypothesis H_0_ stating that the slope of the linear regression is equal to 0.

For both tests, their output is assessed with the computation of the p-value, denoting the probability that the studied sample is observed under H_0._ The significance threshold is set at p=0.05 for the rejection of the null hypothesis. Higher but weak values may indicate that with more data points H_0_ might be rejected as well; thus, it will be reported as a *trend* when the p-value ranges between 0.05 and 0.1.

## 3. Results

### 3.1 General observations

FasGa dye enables the characterization of wood structure composition according to tissue type. This provided a rapid means of assessing the temporal and infection-induced variations and differences with respect to the infection methods used. In xylem and phloem structures, cellulosic tissues appear in blue on the sections, particularly within parenchymatous tissues and tyloses. We can notice (see Fig.8, A and B) that structures with a balanced composition of lignin and cellulose are stained in shades of pink and purple; this color is found in fibers and wood rays.

**Figure 8:**
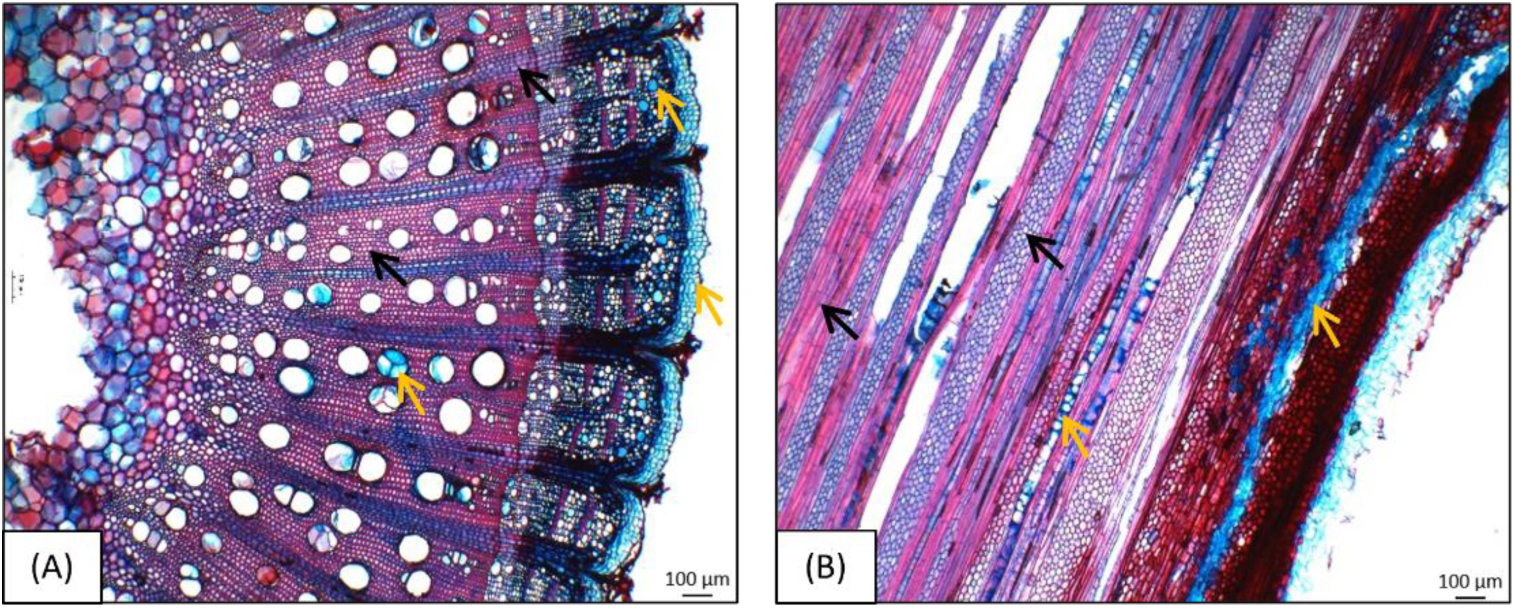
Transverse (A, x4) and tangential (B, x4) sections of healthy Riesling grapevine wood staining with FasGa dye. This coloration highlights cellulose-rich tissues in blue, such as parenchyma cells and tyloses (yellow arrows), as well as lignocellulosic tissues appearing in pink–purple within fibers and rays both in xylem and phloem structures (black arrows).

### 3.2 Absence of temporal and infection variation in vessel morphology within the general xylem

Xylem vessel area differed according to grapevine cultivar and did not seem to vary over time (Fig.9). Gewurztraminer appears to exhibit larger xylem vessels than Riesling, with vessel areas ranging from approximately 4500 to 5000 µm², compared to 3500 to 4000 µm² in Riesling. The different infection treatments do not appear to have any impact on vessel area in either of the studied cultivars (Fig.9).

**Figure 9:**
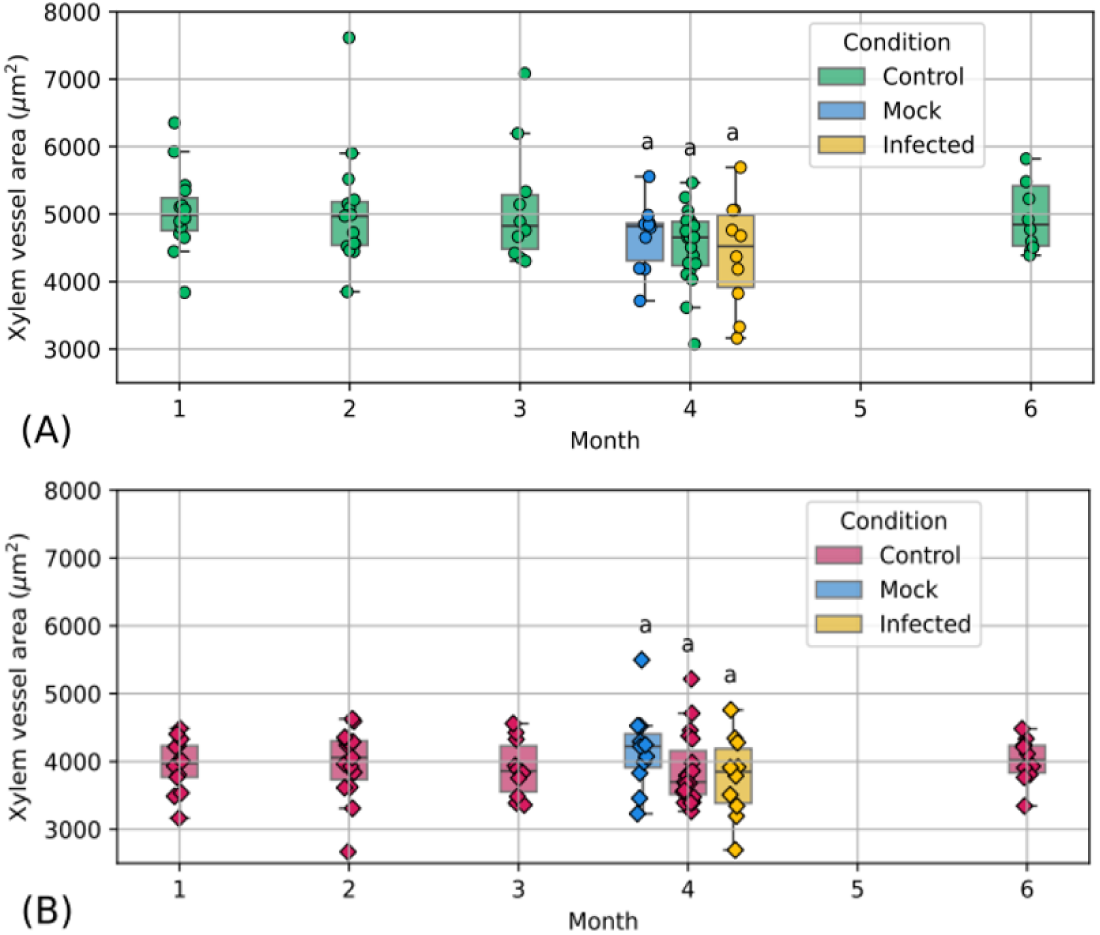
Analysis of xylem vessel area (µm²) across different sampling time points and infection treatments in Gewurztraminer (A) and Riesling (B) cultivars. The analysis was carried out on macroscopic images to provide a comprehensive view of the entire cross section (Fig.4). Here and in the following, sample values are superimposed with the boxplot.

### 3.3 Quantification of dorsoventral vessels repartition in xylem portion

Xylem vessel area, proportion, and eccentricity were measured separately for each time point and for each cultivar, but no significant differences were observed over time (Supplemental Data 1). Because these three parameters seemed month-invariant, data among months were pooled together to compare the two grape varieties with more robustness. Moreover, first observations on cross sections suggested that vessel dimension varies according to their dorsoventral orientation (Fig.4). For this purpose, all samples were oriented along the same dorsoventral axis in order to obtain a representative view of the xylem vessel organization. The vessels were indexed by their position and angle with respect to the dorsoventral axis, and their properties were studied over 32 angular sectors, yielding a polar histogram of the quantities of interest. As previously described, a difference in vessel size between the grape varieties was observed, with a larger surface area for Gewurztraminer than Riesling. However, no variation was observed concerning the frequency and eccentricity of vessels between the two grape varieties (Fig.10).

**Figure 10:**
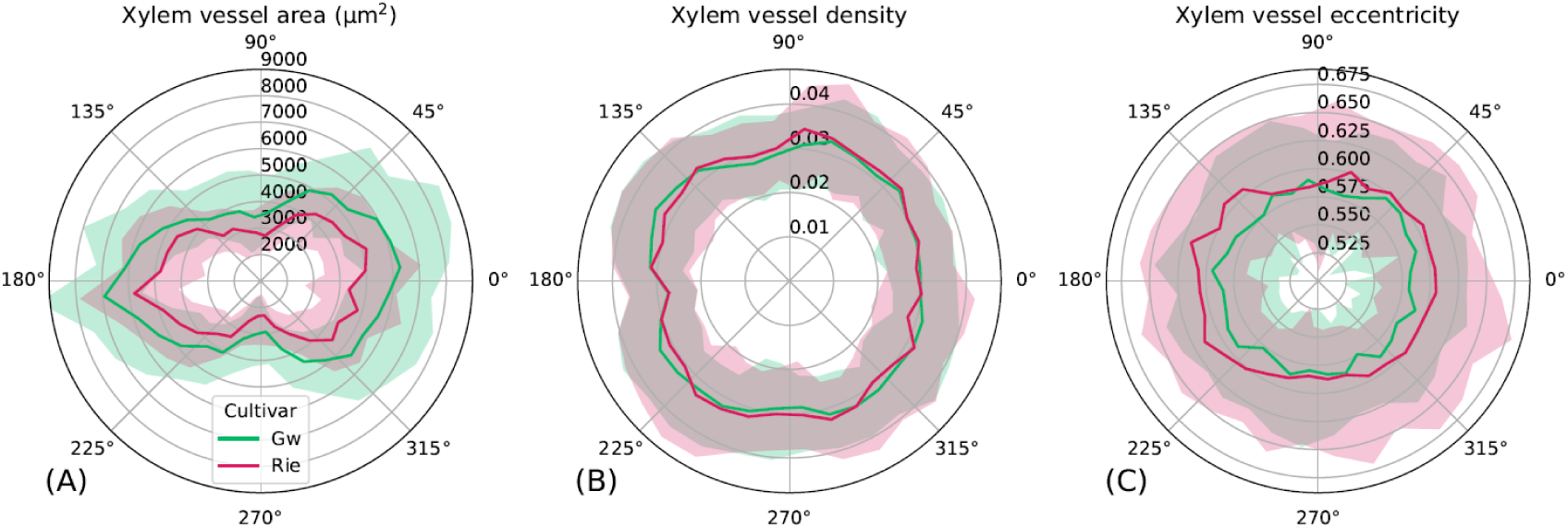
Schematic representation of vessel distribution in cross sections, integrating all sampling time points for each grapevine cultivar. The shaded areas correspond to the interval between the top 10% and 90% of observations, highlighting individual vessel variability. The general xylem organization is illustrated, with quantitative average analyses of vessel area (A), proportion (B), and eccentricity (C). The analysis was carried out on macroscopic images to provide a comprehensive density of the entire cross section.

Interestingly, the dorsoventral arrangements of vessels affect vessel diameter but not vessel density (Fig.10, A-B). Indeed, there is no compensatory increase of vessel number to offset the reduction in sap exchange surface in the dorsoventral or lateral surfaces. Moreover, vessels in plants tended to exhibit similar eccentricity in both cultivars and independently of dorsoventral orientation (Fig.10, C).

The same analysis was carried out on both grapevine cultivars on samples that have undergone treatment: control, mock and infected with *Np*Bt67. We noticed that vessels are organized according to the same dorsoventral plane and that vessel area, frequency, and eccentricity are not affected, regardless of the treatment type (Supplemental Data 2).

### 3.4 Trend toward enhanced new xylem formation during infection

Morphologically, a ring boundary can be observed, separating newly formed xylem from older xylem; this demarcation delimits the ring limit (see Fig.8, A). Measurements show that there was no clear month-wise tendency concerning the development of new xylem thickness, excepting for month 1 in Riesling cultivar, which we discarded because many samples had low or no new xylem. Gewurztra-miner appears to form a wider newly formed xylem than Riesling. This phenomenon does not increase over time and remains stable. No significant variation in xylem thickness was detected over time in untreated plants regarding the two grapevine cultivars (Fig.11). In contrast, in month 4, plants subjected to drilling and inoculation exhibited a tendency toward increased thickness of newly formed xylem tissue. A statistically significant increase was observed in Gewurztraminer plants infected with the *Np*Bt67 strain (p = 8,10^-4^) compared to the non-inoculated and non-drilled control. For Riesling, the drilling treatment alone influences the thickness of the newly formed xylem (p = 0.033), while no clear effect could be attributed to pathogen infection even though they appear to follow the same trend according to the statistical results (p = 0.07) (Fig.11).

**Figure 11:**
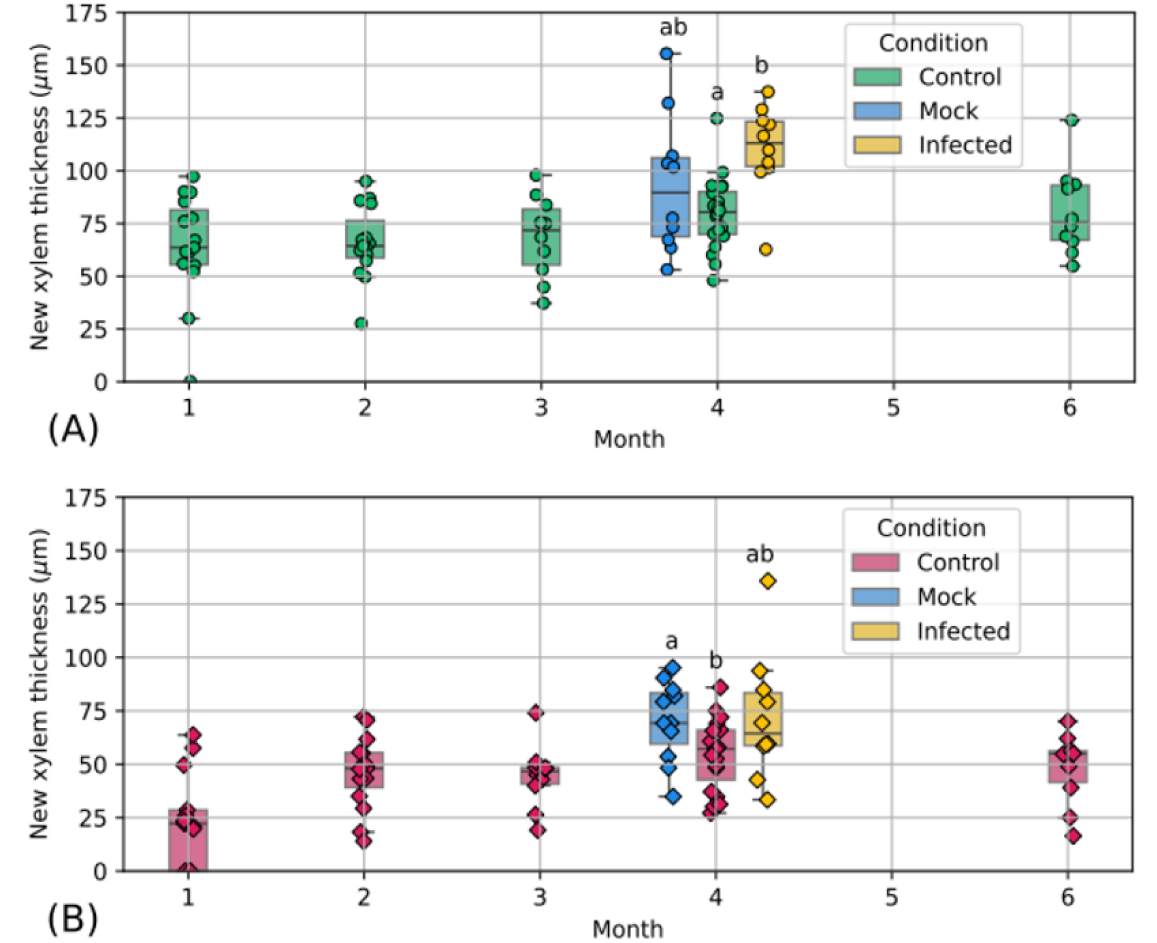
Analysis of newly formed xylem thickness across different sampling time points and infection treatments in Gewurztraminer (A) and Riesling (B) cultivars. The analysis was carried out on macroscopic images to provide a comprehensive view of the entire cross section.

### 3.5 New xylem shows a temporal increase in vessel number, with no infection-related effects on vessel formation or vessel area

A significant increase in the number of vessels formed in the new xylem was observed over time in Gewurztraminer (p = 0.016). This trend appears to be similar in Riesling (p = 0.057). The different treatment conditions applied to the plants did not seem to affect the formation of new vessels in this developing tissue (Fig.12).

**Figure 12:**
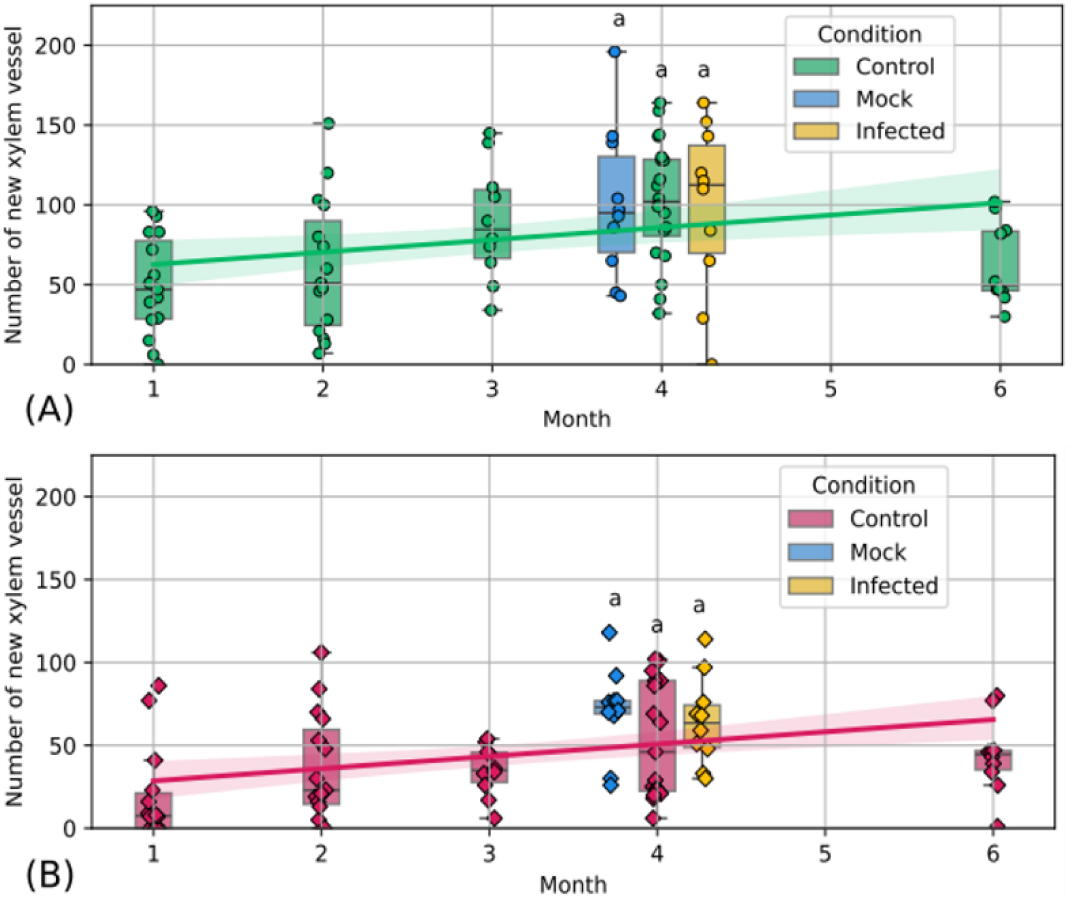
Analysis of number of vessels of newly formed xylem across different sampling time points and infection treatments in Gewurztraminer (A) and Riesling (B) cultivars. The analysis was carried out on macroscopic images to provide a comprehensive view of the entire cross section. Here and in the following, the linear regression is represented when the corresponding p-value is below 0.1. Shaded areas correspond to the 95% confidence interval, estimated over 10^4^ bootstrap resamples.

Samples were quite heterogeneous with regard to the area of newly formed vessels. No temporal variation in vessel area was observed in this developing tissue. Infection treatments had no effect on Gewurztraminer; only a significant difference was observed in Riesling between the undrilled control and the mock-drilled sample inoculated with PDA (p = 0.01) (Fig.13). In this case, an increase in the area of newly formed vessels was observed in Riesling, but this appears to be independent of fungal infection.

**Figure 13:**
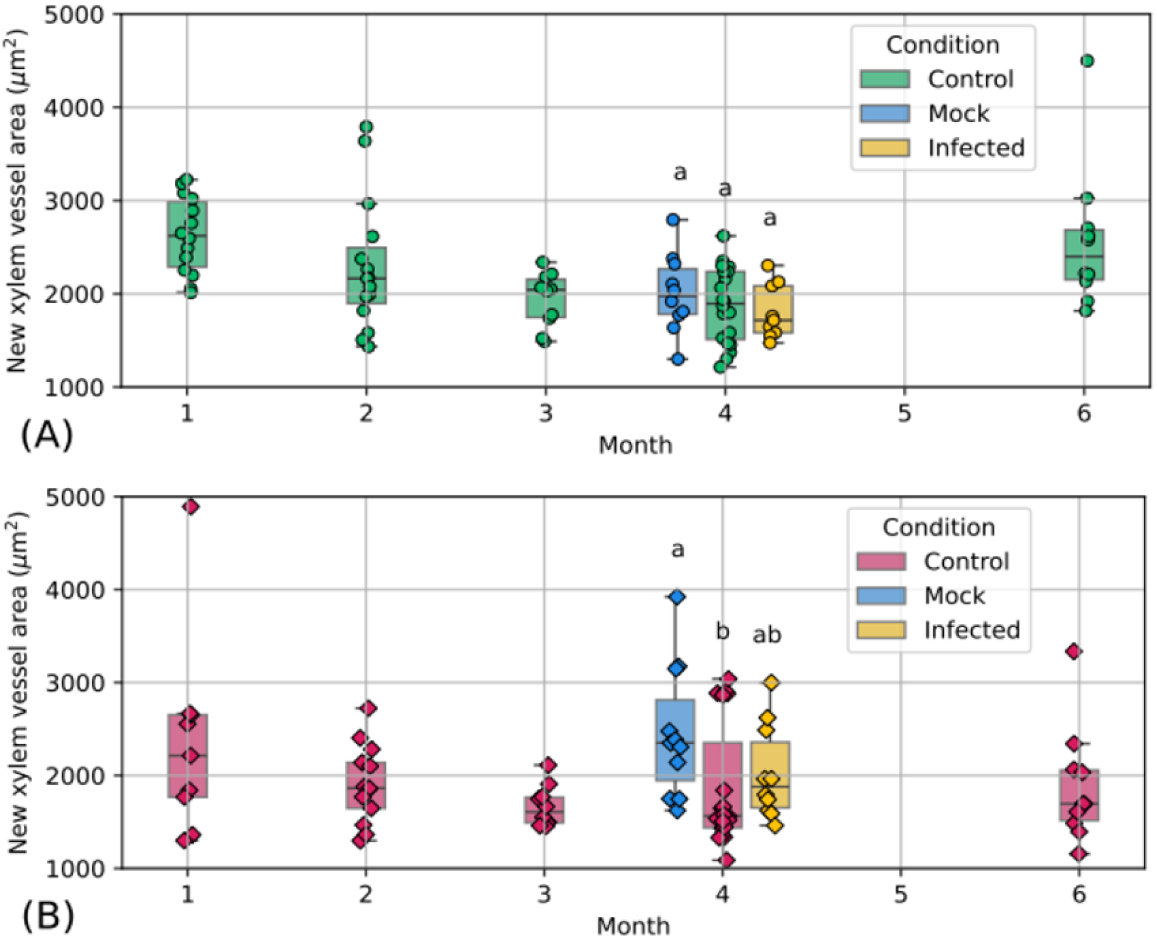
Analysis of new xylem vessel area (µm^2^) across different sampling time points and infection treatments in Gewurztraminer (A) and Riesling (B) cultivars. The analysis was carried out on macroscopic images to provide a comprehensive view of the entire cross section.

### 3.6 Stress-associated lignification accompanies temporal cellulosic tissue formation

A progressive reduction in red staining intensity was detected over time, while blue coloration increased correspondingly. This pattern is significant for Riesling (p_blue_ = 6.10^-4^, p_red_ = 6.10^-4^) and tends to the same observation for Gewurztraminer (p_blue_ = 0.07, p_red_ = 0.0507). Regarding the effect of the different treatment modalities, a significant increase in red staining was observed in GW samples that were drilled and inoculated with either PDA (p_mock/control_ = 0.01) or *Np*Bt67 (p_infected/control_ = 10^-3^) compared to the control. However, this variation appears to be more associated with the drilling itself than with the fungal effect because no significant difference was found between samples inoculated with PDA and *Np*Bt67. A similar trend was observed in Riesling (p_mock/control_ = 0.07 and p_infected/control_ = 0.06). In parallel, no significant variation in blue staining intensity was observed with respect to treatment modality or cultivar (Fig.14). It would appear that the wood tissue lignified when subjected to stress. Nevertheless, the coloration protocol associated with the image analysis pipeline allowed to quantify grapevine’s response to wounding damage in woody tissues.

**Figure 14:**
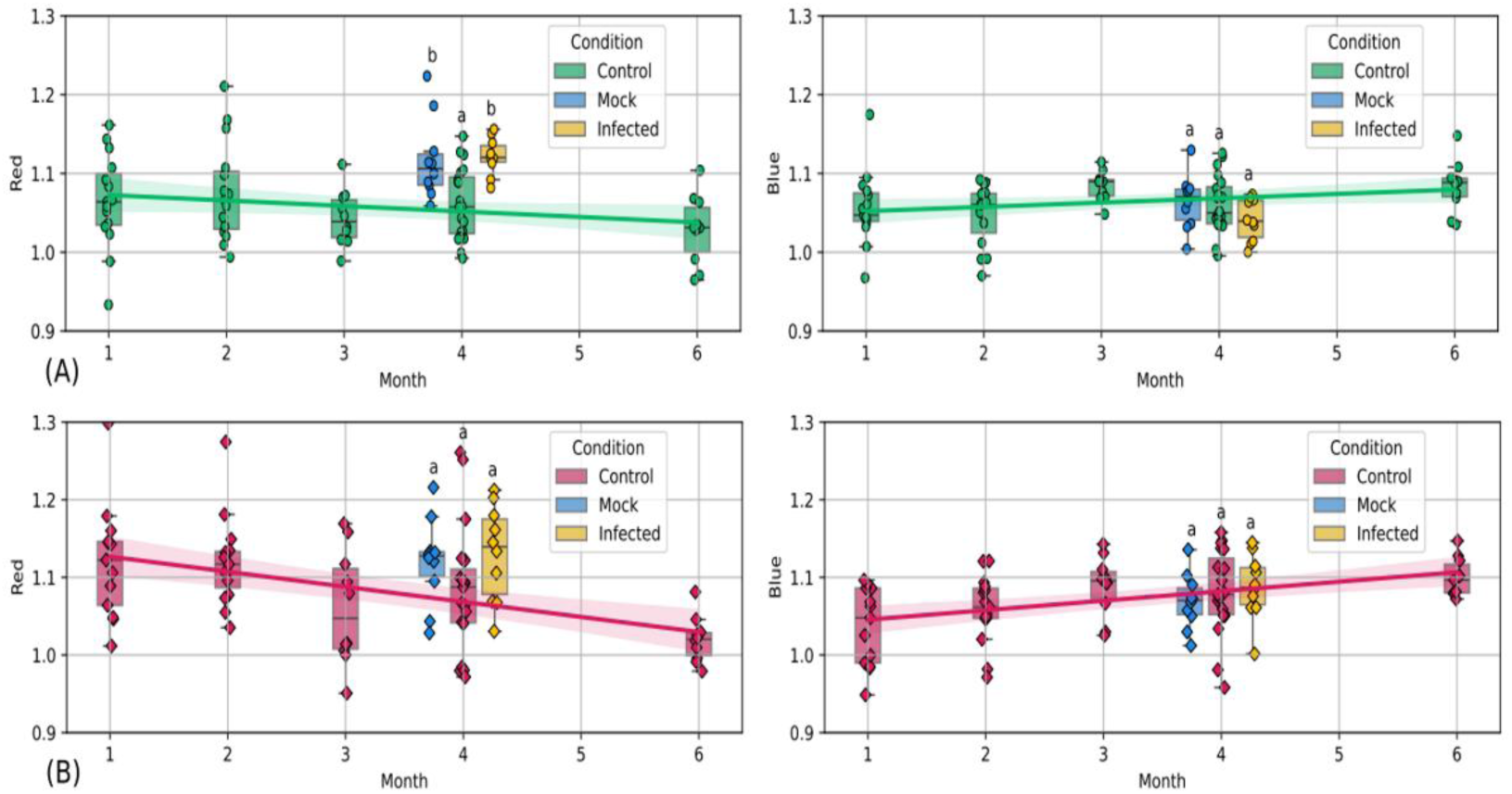
Analysis of overall coloration on cross sections across different sampling time points and infection treatments in Gewurztraminer (A) and Riesling (B) cultivars. Red (left) and blue (right) intensity levels were monitored on each section. The analysis was performed on microscopic x4 images of cross-section slices.

### 3.7 Phloem structure shows no temporal variation but exhibits cultivar-dependent predisposition

The phloem was examined in the radial section to facilitate the measurement of its thickness. In these conditions, no significant variation of phloem thickness was observed over time and between the studied cultivars. Additionally, the different treatments applied to plants did not appear to affect phloem growth. However, the phloem tissue seems to be thicker in Riesling than in Gewurztraminer (Fig.15).

**Figure 15:**
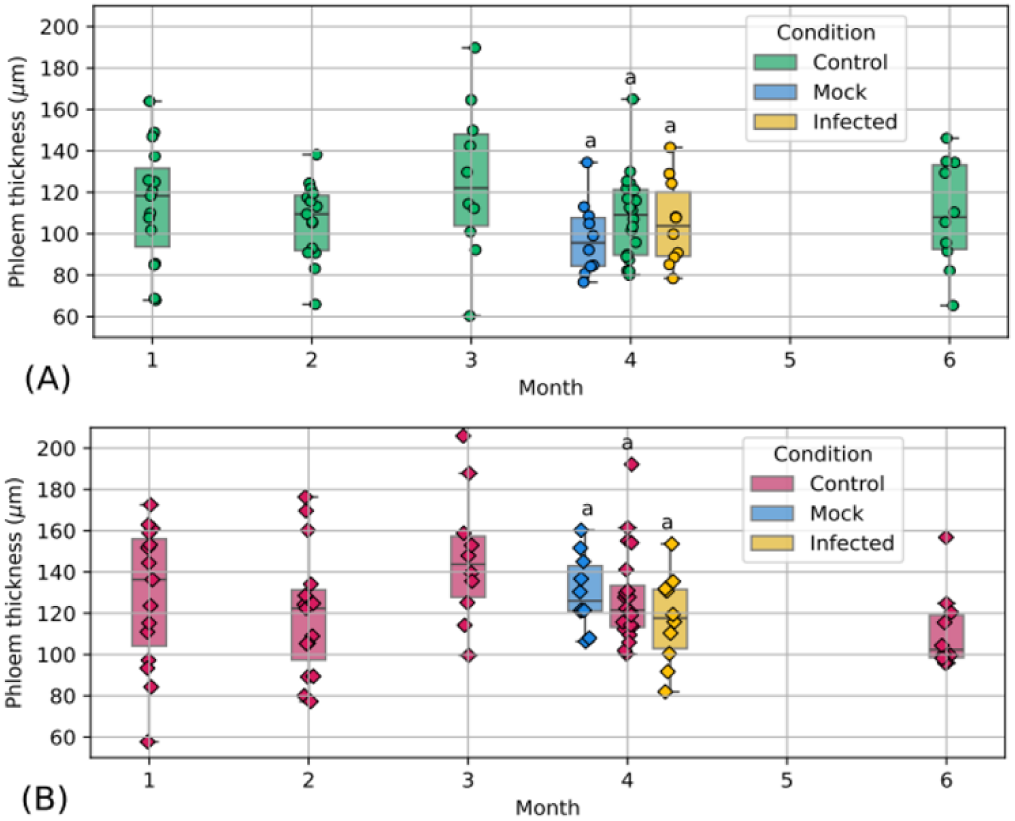
Analysis of phloem thickness across different sampling time points and infection treatments in Gewurztraminer (A) and Riesling (B) cultivars. Analysis was conducted on microscopic radial sections at x4 magnification.

### 3.8 Observation of wood discoloration in the absence of infection

No temporal variation in wood composition was macroscopically observed during these assays as described above, except in regions exhibiting discoloration. Indeed, spontaneous internal discoloration could be observed in several samples in the absence of any experimentally induced infection. In this case, lignified tissues are observed on fractions presenting discoloration. Indeed, a more intense red staining with FasGa was detected in samples exhibiting discoloration. We found that 33% of the Gewurztraminer samples and 35% of the Riesling samples exhibited discolorations. These discolorations were clearly defined and macroscopically visible during sampling. In microscopy, FasGa staining also confirmed the presence of discoloration, as lignified wood structures stained in red, consistent with tissue lignification/suberization (Fig.16). The same phenomenon was observed in control samples from the plants used to assess wood variation to fungal infection. In these cases, discolorations were more marked, as 60% of Gewurztraminer and 100% of Riesling samples exhibited discoloration.

**Figure 16:**
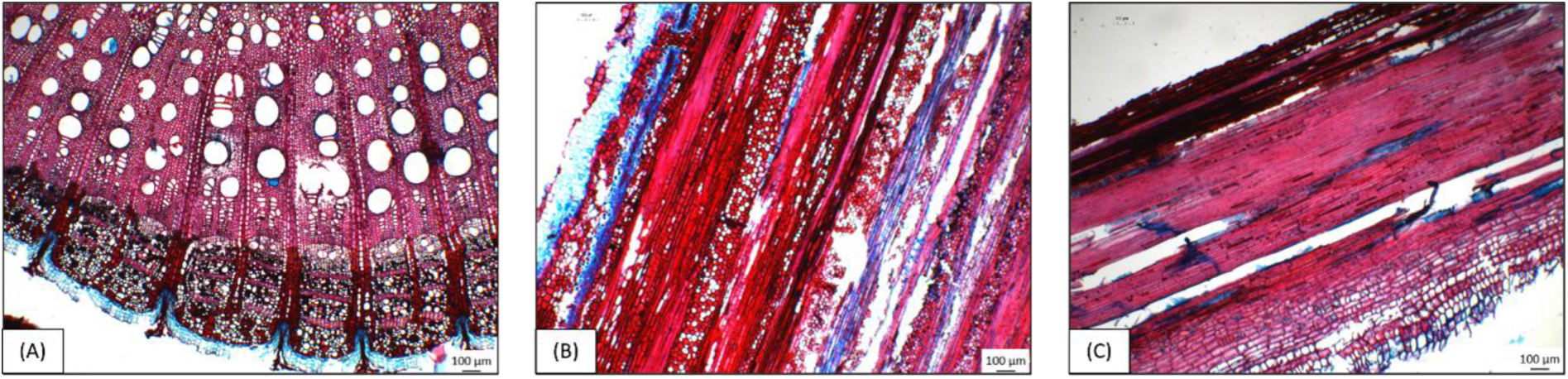
Tissues presenting wood discoloration in transverse (A, x4), tangential (B, x4) or radial (C, x4) sections appears predominantly stained in red with FasGa Dye with predominantly lignified tissues.

This discoloration-specific feature led us to assess its impact on the wood anatomy investigated in this study. Data were compared with samples showing no discoloration across all parameters subsequently analyzed for the two cultivars (Supplemental Data 3). Structural analyses of the wood revealed no detectable differences associated with discoloration. Samples exhibiting internal discoloration in the absence of induced infection were thus included in the study and considered as ’healthy/normal’ samples for the rest of the study.

## 4. Discussion

### 4.1 General xylem vessels morphology

In this study, the temporal dynamics of wood structure were examined in two grapevine cultivars susceptible to trunk diseases: Riesling and Gewurztraminer. No significant temporal changes were detected in xylem morphology, including vessels areas, vessels number, and eccentricity for each cultivar. This portion of the wood, which was already formed and present at the beginning of the experiment, did not exhibit any temporal changes.

However, results revealed a specific, distinct dorsoventral spatial organization characteristic of grapevine wood [39]. Image processing analysis showed that this organization did not affect vessels number, which remained consistent across all regions of the cross section. In contrast, vessels area decreased in the lateral regions corresponding to the bud side, indicating localized vessel narrowing that may restrict sap flow in this zone. The presence of vessels with smaller diameters may help reduce the risk of embolism and vessel blockage caused by pathogenic fungi [32]. Moreover, since this distribution pattern was observed in both cultivars, it suggests that the phenomenon is inherent to grapevine anatomy rather than cultivar-dependent. However, vessel areas do vary between cultivars, and this variation is associated with the tolerance of grapevine cultivars to GTDs. In this study, the fact that Gewurztraminer exhibits larger vessels than Riesling is consistent with its higher susceptibility to trunk diseases. Several studies investigating xylem anatomy have demonstrated cultivar-dependent variations in vessel diameter, which have been linked to differences in susceptibility to grapevine trunk diseases [33,40].

Vessel eccentricity alone does not appear to constitute an independent resistance-related trait. Moreover, there is, to our knowledge, limited data available on this aspect in literature, highlighting the need for additional experiments on other cultivars to confirm its potential role as an aggravating factor in sensibility to GTDs.

### 4.2 New xylem: thickness, area and vessels frequency

When focusing on the recently formed xylem zone, a greater development of this tissue was observed in Gewurztraminer compared to Riesling. This difference could be partly explained by the initial cane diameter for each grapevine cultivar used for the experiment. Gewurztraminer plants indeed exhibited thicker and more vigorous canes than Riesling. At the start of the experimentation, this parameter may have influenced radial growth and should therefore be considered when interpreting these results. To our knowledge, there is no data currently available in the literature on this aspect, which highlights the need to repeat the experiment to take into account the uniformity of canes for both cultivars.

Furthermore, in this study no significant variation in the thickness of the newly formed xylem was observed over time in both grapevine cultivars. The absence of observable changes in xylem thickness over time may be attributed to the short duration of the experiment and the stable environmental conditions, which together may have constrained secondary growth.

Interestingly, our results suggest that infection by *Np*Bt67 may induce enhanced development of the newly formed xylem, while no effect was observed on old xylem general anatomy. This observation was particularly evident in Gewurztraminer, although Riesling appeared to follow a similar trend. This pattern coincides with compartmentalization of the pathogen in plants, which seems to stimulate the formation of new tissue to compensate for the vessel colonization by fungi while maintaining the sap flow. A similar response was also observed following mechanical wounding caused by drilling of the plants, which appears to trigger increased formation of new xylem tissue. These observations represent preliminary trends and require further replication to be confirmed. Nonetheless, the results suggest that mechanical injury combined with pathogen inoculation may promote enhanced formation of secondary xylem in Gewurztraminer cultivar. Finally, the infection treatments applied to the plants did not appear to affect vessel area. While new xylem tissue was produced, there was no compensatory change in vessel diameter.

Note that a study reported a reduction in xylem formation in Chardonnay, with narrower growth ring boundaries in plants subjected to both biotic and abiotic stresses, such as water deficit and infection by Flavescence dorée [29]. In this case, it appears that the plant restricts its growth and the formation of new tissue in order to conserve energy and reduce sap flow. The plant thus seems to modulate its developmental processes according to the type and intensity of stress encountered. Besides, this phenomenon is cultivar-dependent. On the other hand, a change in cambial vascular activity, was observed by Dell’Acqua and al. in vines exhibiting Esca symptoms with the presence of several tyloses in xylem. In this case, the plant compensates by producing a large number of new smaller vessels, thereby maintaining sap conduction within the stem [41].

### 4.3 Colorimetry analysis

Monitoring the coloration of our transverse sections allowed us to observe changes in wood structure composition over time. A gradual decrease in red intensity was observed, coinciding with an increase in blue staining. These results are consistent with the formation of new tissue rich in cellulose, as lignification had not yet been completed. Image analysis also revealed tissue suberization when plants were wounded or wounded and infected by *Np*Bt67. Consequently, this tool enables straight-forward, automated quantification of color changes associated with plant response to environmental stress, wounding damage in the present work. As previously demonstrated by Pouzoulet *et al.*, infection triggers localized responses, including suberization of the cell walls and also lignification of affected tissues [33].

### 4.4 Phloem thickness

The analysis of the phloem did not reveal any anatomical variation over time or in response to drilling or infection. Phloem tissue is less developed than xylem tissue over time and likely requires a longer developmental period to detect any potential effect, if such effect exists. To our knowledge, this portion of the stem has received little attention in the literature compared with the xylem, preventing any direct comparison with previously published data. Nevertheless, our results indicate that phloem thickness appears to be greater in Riesling than in Gewurztraminer. This increase may favor the resilience of the plant and remain to be correlated to the tolerance observed in vineyard conditions [13, 31].

### 4.5 Wood discoloration did not impact anatomical variation in wood anatomy

During these experiments, wood discoloration was observed in plant samples that had not undergone any treatment (drilling or inoculation). This phenomenon had not previously been reported in our laboratory. Although this phenomenon did not affect the wood anatomy examined in this study, it is an important parameter to consider for future studies, depending on the objectives of the investigation. The image analysis pipeline enabled to isolate the impact of fungal infection from those external factors that probably caused these discolorations.

## 5. Conclusion

This study investigated wood anatomy in two Alsatian cultivars susceptible to GTDs. Although high inter-plant variability and contrasting responses within our experimental model preclude definitive conclusions, our results suggest a potential rearrangement of newly formed xylem anatomy during infection. This phenomenon remains to be confirmed through further experiments involving longer cultivation periods and additional susceptible cultivars. We also validated a robust FasGa staining protocol for reliable visualization of wood structures. This coloration, together with the image analysis pipeline, allowed to quantify tissue’s lignification in response to wounding damage. In addition, the computational model developed here provides a powerful and reliable analytical tool for high-throughput sample processing and for the quantitative assessment of anatomical traits that have remained largely unexplored.

Finally, all codes and microscopy images generated in this study have been made publicly available, establishing a comprehensive and accessible resource for the scientific community, to support ongoing hypotheses. If further studies could investigate anatomical variations of woody tissues over longer time periods, this tool could also be adapted to microscopy samples from older plants in vineyard conditions.

## Acknowledgement

This research was funded by the Université de Haute-Alsace research grant Vineside. Authors are grateful to Julien RUELLE (Silvatech, INRAE Champenoux) for training in sectioning and staining techniques. They thank Jacky MISBACH (UEAV, https://doi.org/10.15454/1.5483269027345498E12, INRAE Colmar) for providing plant material and Catherine REINBOLD (UMR 1131 SVQV, INRAe Colmar) for access to the microscopy platform, where macroscope observations were performed. The Plant Innovation Platform (UR 3991, LVBE-UHA) is acknowledged for plant cultivation and experimental support. The authors further thank Olivier HAEBERLE and Bruno COLICCHIO (UR 7499, IRIMAS-UHA) for complementary fluorescence microscopy images acquisition, which was awarded the Optica OPN image of the week (week of February 20, 2026).

## Data availability

All microscopy images can be accessed on the Zenodo platform [35] at 10.5281/ze-nodo.18850060. The code enabling automated microscopy image analysis is available online at github.com/courbot/vineside.

## Authors contributions

CP, JBC, YL and RJGP conceived and designed the experiments;

CP, YL and RJGP performed the experiments;

CP performed sample preparation and microscopy;

CP, JBC, and RJGP analyzed and interpreted the data;

CP, JBC, YL and RJGP wrote the paper.

---

**Supplemental Data 1:**
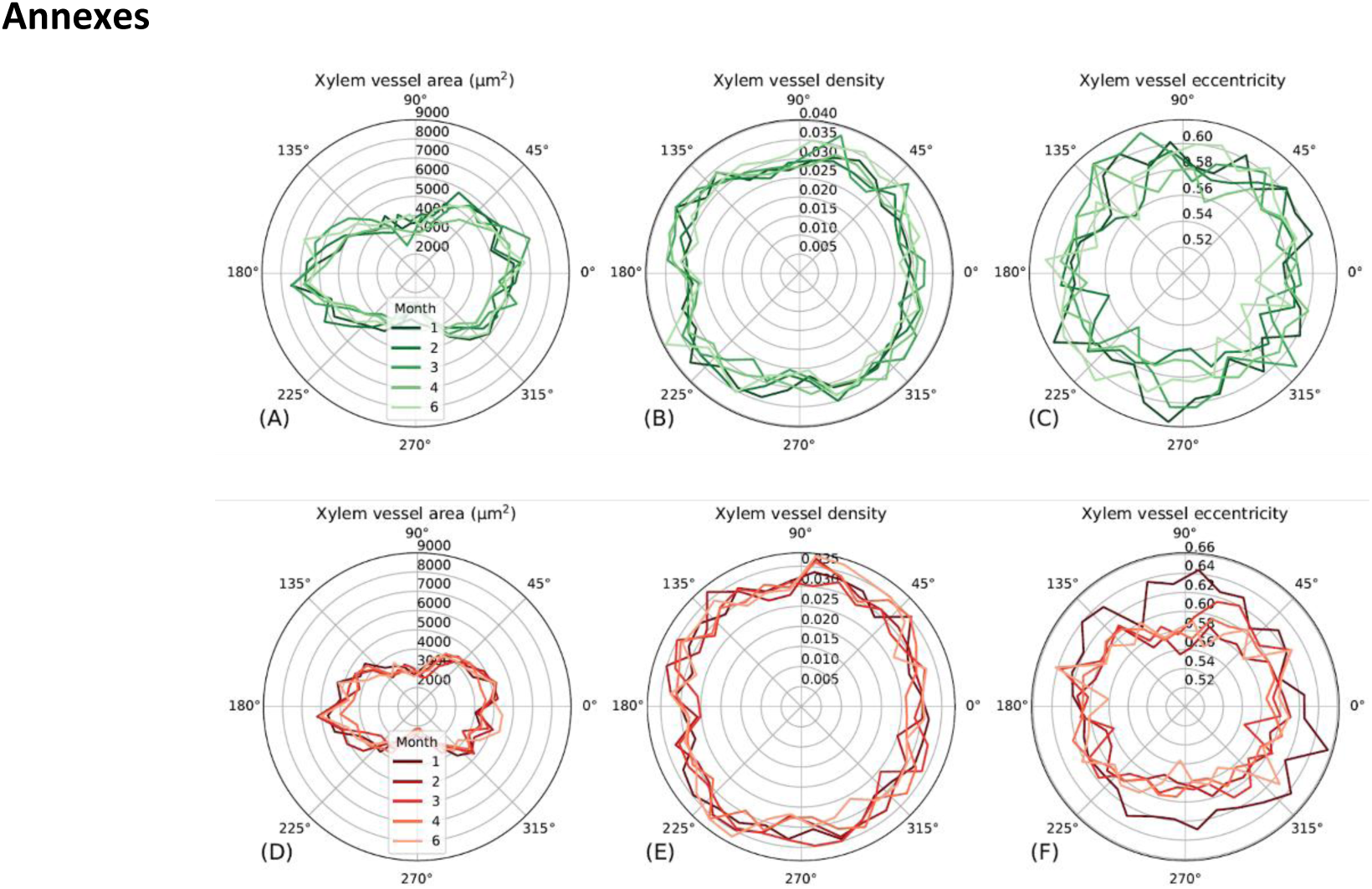
Schematic representation of xylem vessel distribution in cross sections, at each sampling time point in Gewurztraminer (A-C) and Riesling (D-F). The general xylem organization is illustrated, with quantitative analyses of vessel area (A, D), proportion (B, E), and eccentricity (C, F). The analysis was carried out on macroscopic images to provide a comprehensive view of the entire cross section.

**Supplemental Data 2:**
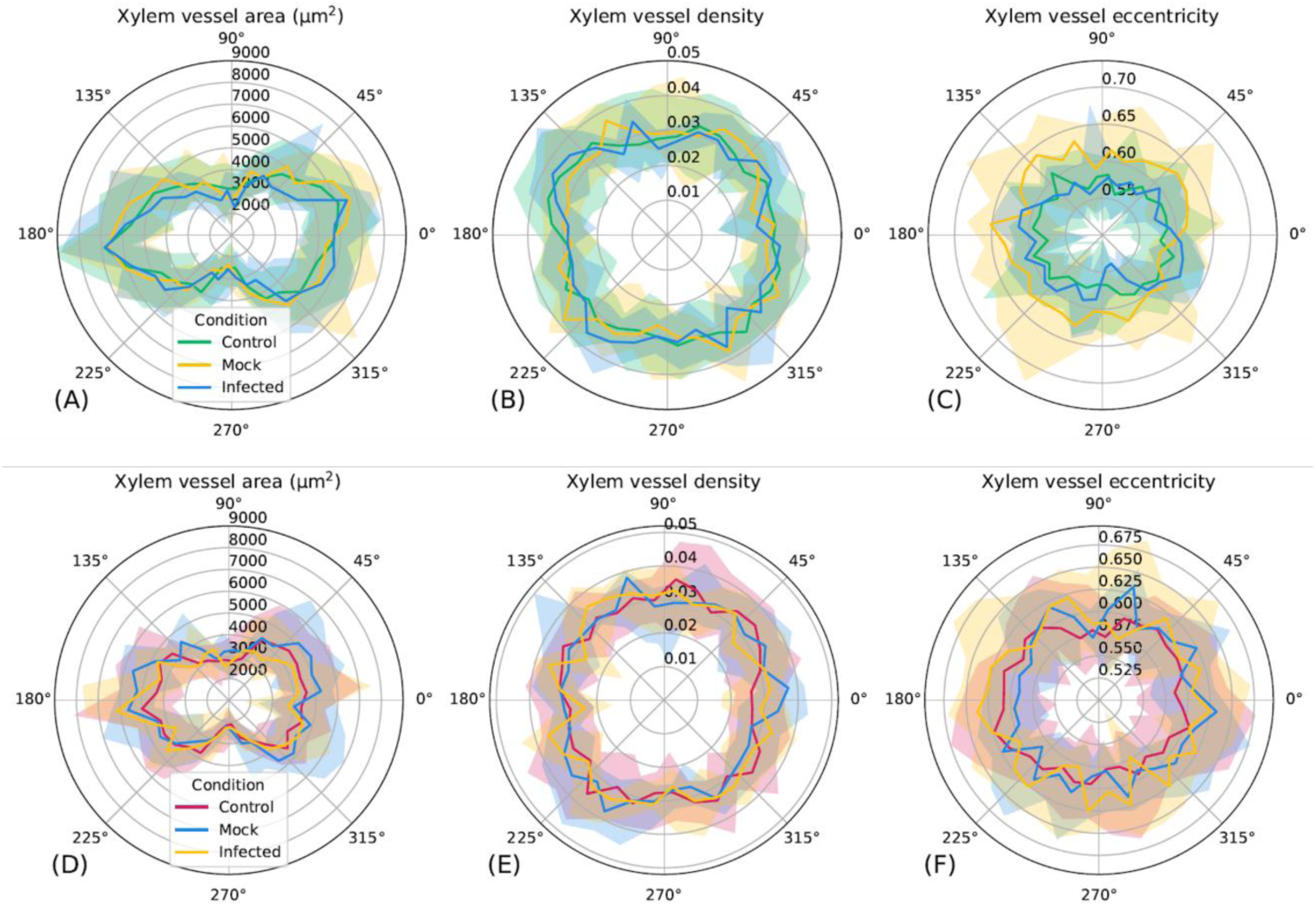
Schematic representation of the general xylem organization in two grapevine cultivars: Gewurztraminer (A-C) and Riesling (D-F), following control, mock or NpBt67 infection treatments. Quantitative analysis was conducted on vessel area (A, D), proportion (B, E) and eccentricity (C, F).

**Supplemental Data 3:**
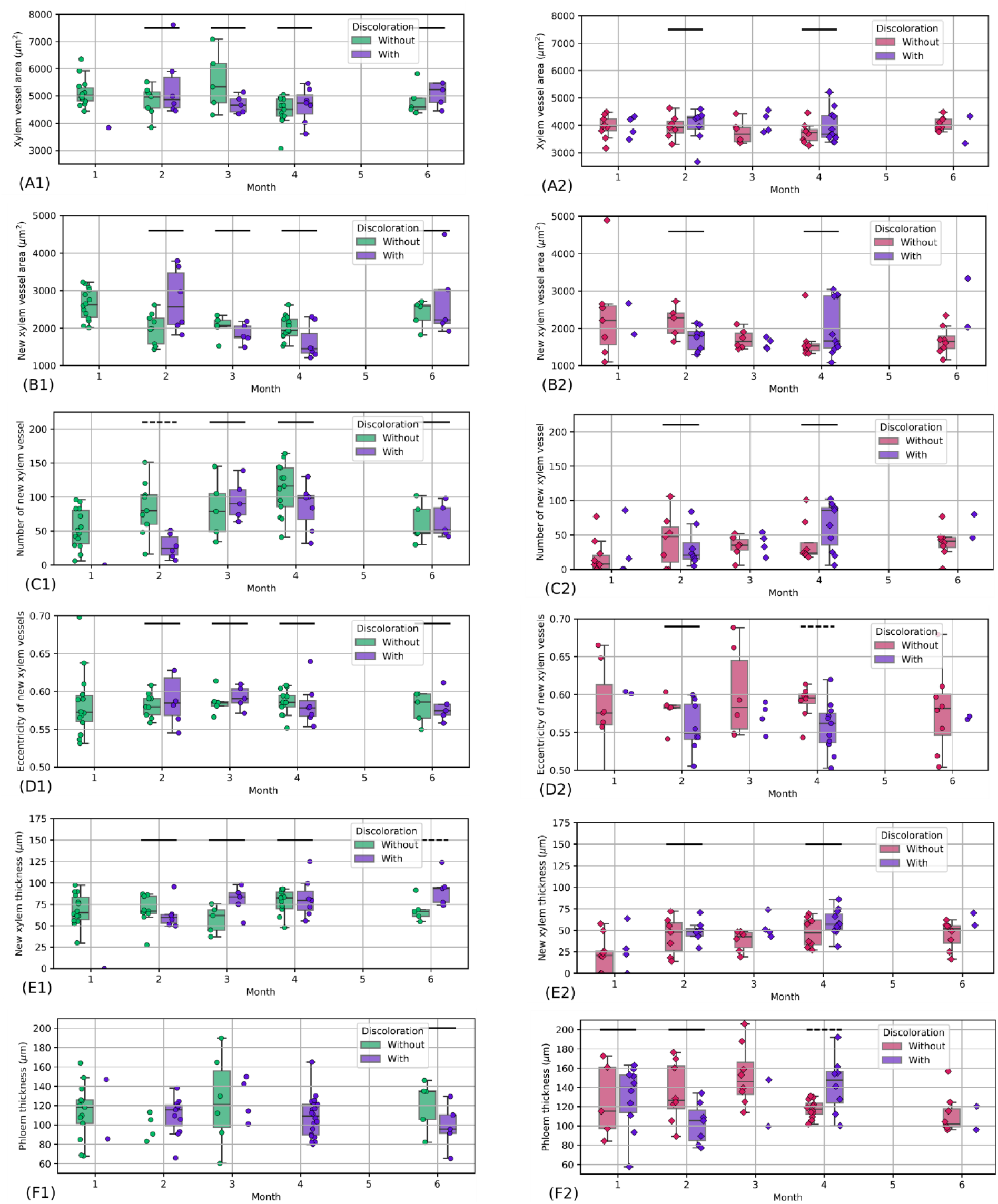
Investigation of the effects of wood discoloration on xylem and phloem anatomical traits, including total xylem vessel area (A); new xylem vessel area (B), density (C) and eccentricity (D); thickness of newly formed xylem ring (E); and phloem thickness (F) in two grapevine cultivars, Gewurztraminer (1) and Riesling (2). Significant differences between discolored and non-discolored tissues are indicated by horizontal dashed bars, whereas non-significant differences are shown with solid bars. Note that for small samples (< 5 values) the boxplot is not displayed and the difference is not tested. Only 4 comparisons between samples with and without discoloration showed significant differences across the 34 total times steps analyzed.

